# Conserved tandem arginines for PbgA/YejM allow *Salmonella* to regulate LpxC and control lipopolysaccharide biogenesis during infection

**DOI:** 10.1101/2020.12.07.415661

**Authors:** Nicole P. Giordano, Joshua A. Mettlach, Zachary D. Dalebroux

## Abstract

*Salmonella enterica* serovar Typhimurium uses PbgA/YejM, a conserved multi-pass transmembrane protein with a soluble periplasmic domain (PD), to balance the glycerophospholipid (GPL) and lipopolysaccharide (LPS) concentrations within the outer membrane (OM). The lipid homeostasis and virulence defects of *pbgAΔ191-586* mutants, which are deleted for the PD, can be suppressed by substitutions in three LPS regulators, LapB/YciM, FtsH, and LpxC. We reasoned that *S*. Typhimurium uses the PbgA PD to regulate LpxC through functional interactions with LapB and FtsH. In the stationary phase of growth, *pbgAΔ191-586* mutants accumulated LpxC and overproduced LPS precursors, known as lipid A-core molecules. Trans-complementation fully decreased the LpxC and lipid A-core levels for the mutants, while substitutions in LapB, FtsH, and LpxC variably reduced the concentrations. PbgA binds lipid A-core, in part, using dual arginines, R215 and R216, which are located near the plasma membrane. Neutral, conservative, and non-conservative substitutions were engineered at these positions to test whether the side-chain charges for residues 215 and 216 influenced LpxC regulation. Salmonellae that expressed PbgA with dual alanines or aspartic acids overproduced LpxC, accumulated lipid A-core and short-LPS molecules, and were severely attenuated in mice. Bacteria that expressed PbgA with tandem lysines were fully virulent in mice and yielded LpxC and lipid A-core levels that were similar to the wild type. Thus, *S.* Typhimurium uses the cationic charge of PbgA R215 and R216 to down-regulate LpxC and decrease lipid A-core biosynthesis in response to host stress and this regulatory mechanism enhances their virulence during bacteremia.

**IMPORTANCE:** *Salmonella enterica* serovar Typhimurium causes self-limiting gastroenteritis in healthy individuals and severe systemic disease in immunocompromised humans. The pathogen manipulates the immune system of its host by regulating the lipid, protein, and polysaccharide content of the outer membrane (OM) bilayer. Lipopolysaccharides (LPS) comprise the external leaflet of the OM, and are essential for establishing the OM barrier and providing gram-negative microbes with intrinsic antimicrobial resistance. LPS molecules are potent endotoxins and immunomodulatory ligands that bind host-pattern receptors, which control host resistance and adaptation during infection. Salmonellae use the cationic charge of dual arginines for PbgA/YejM to negatively regulate LPS biosynthesis. The mechanism involves PbgA binding to an LPS precursor and activating a conserved multi-protein signal transduction network that cues LpxC proteolysis, the rate-limiting enzyme. The cationic charge of the tandem arginines is critical for the ability of salmonellae to survive intracellularly and to cause systemic disease in mice.

## INTRODUCTION

*Salmonella enterica* serovar Typhimurium (*S*. Typhimurium) infects humans that have ingested contaminated food or water, or that have interacted with companion animals or livestock (1,2). The resulting gastroenteritis is highly inflammatory but mostly self-limiting and rarely requires therapeutics. Immunocompromised humans are susceptible to non-typhoidal bacteremia, a systemic disease that occurs when pathogens penetrate the mucosal and epithelial barriers of the intestine, enter the lymphatic system, and eventually inhabit the bloodstream (1). During systemic infection, *S*. Typhimurium largely survives as a facultative intracellular pathogen and manipulates the immune system of its host from within the endocytic vacuoles of macrophages and dendritic cells (3). The acidic pH and environment of the late endosome causes *S.* Typhimurium to increase the activity of a multitude of regulatory proteins and mechanisms, many of which control the glycerophospholipid (GPL) and lipopolysaccharide (LPS) content of the outer membrane (OM) (4). Maintaining and regulating OM-lipid content is critical to nearly every aspect of *S.* Typhimurium pathogenesis and contributes to intrinsic antimicrobial resistance (4–6).

The OM is an asymmetrical bilayer of GPLs in the inner leaflet and lipopolysaccharides (LPS) in the outer leaflet (7–11). The asymmetric character of the OM allows to the peripheral bilayer to function as a physiochemical barrier, which promotes defensive functions (12, 13). Lipid A molecules are multi-acylated, phosphorylated disaccharolipids that comprise the OM outer leaflet. Lipid A is the amphipathic component of LPS that directly interacts with GPLs (12, 13). Divalent cations form salt-bridges between adjacent phosphates on lipid A molecules to provide lateral stability to the surface, and hydrophobic interactions between the acyl chains of LPS and GPL molecules establish the barrier (12, 14, 15). The biochemical properties of the OM enhance enterobacterial resistance to small hydrophobic antibiotics and promote virulence and disease pathogenesis (4, 7, 13, 16–18).

Enterobacteriaceae use the Lpx, Kdt, and Waa/Rfa enzymes to independently synthesize the lipid-A disaccharolipids and the core oligosaccharides in the cytosol. Next, lipid A and core oligosaccharide are assembled into lipid A-core, the principal LPS precursor. This occurs on the cytoplasmic leaflet of the plasma membrane (13, 19–22). MsbA flips lipid A-core molecules into the periplasmic leaflet of the IM where WaaL/RfaL ligates lipid A-core to the O-polysaccharides (also known as O-antigens) (13, 21, 22). Enteropathogenic *E. coli* and *S. enterica* produce LPS molecules that are decorated with O-antigens of varying polysaccharide chain length (5, 16, 21, 23). The O-antigens are synthesized in the cytosol and attached to undecaprenyl phosphate (Und-P) carrier lipids at the inner leaflet of the IM. Und-P-linked O-antigen conjugates are flipped into the periplasmic leaflet of the IM and polymerized (13, 24). *S.* Typhimurium uses at least two polymerases to produce three O-antigen LPS subtypes, the short (2-15 repeating units; RU), long (16-35 RU), and very long (>100 RU) LPS modalities (4, 24). The LPS structures are transported outward across the periplasm and inserted into the outer leaflet of the OM by the Lpt machinery (20–22). Lipid A-core molecules that are devoid of the O-antigen are also transported to the OM and are intrinsic components of the outer leaflet; thus, O-antigen attachment is not a prerequisite for transport (5, 13, 22).

During stress, enterobacteriaceae regulate wholesale production of lipid A-core and LPS molecules by decreasing the rate of biosynthesis. This occurs through a proteolytic mechanism, which involves LapB/YciM, FtsH, and LpxC (16, 25). LapB is an IM-tethered cytosolic protein whose expression and activity levels increase during stress (25–29). LpxC is a cytosolic deacetylase that catalyzes a rate-limiting step for lipid A-core biosynthesis. Activated LapB molecules prompt the IM protease, FtsH, to degrade LpxC; however, the signals that control LapB and FtsH activity on LpxC are poorly understood (13, 16, 19). *E. coli* LapB binds both LpxC and FtsH and influences LpxC stability, but the specificity of the LapB-LpxC and LapB-FtsH interactions are not known (27, 30). Current literature classifies LapB and FtsH as negative regulators of enterobacterial LPS biosynthesis (27–29).

Work from our lab and others unveiled a fourth component of this regulatory network, PbgA/YejM (5, 31–34). PbgA is an essential IM protein with a large non-essential periplasmic domain (PD) (5, 35). *S*. Typhimurium relies on the PD of PbgA to control the levels of lipid A-core on the OM, in part, through functional interactions with LapB, FtsH, and LpxC (5, 35). The *S*. Typhimurium and *E. coli* PbgA and LapB proteins exhibit a high degree of sequence identity (88.23% and 93.4% identity, respectively). *E. coli* PbgA interacts with LapB through the transmembrane (TM) domains. The essential function of PbgA in laboratory *E. coli* K-12 involves stabilizing LpxC; however these domestic microbes produce a lipid A-core glycolipid on their OM outer leaflet that lacks the O-antigen (5, 31–34). High-resolution crystal structures of the *S*. Typhimurium and *E. coli* PbgA proteins revealed a lipid A-core molecule bound, in part, via tandem arginines, R215 R216, in the non-essential basic region of the PD (34, 35). In a previous study, we demonstrated that *S.* Typhimurium uses the PbgA R215 R216 to enhance the OM-barrier, but we had not tested the role of these residues in regulating LpxC and lipid A-core (35). We interrogated the hypothesis that *S*. Typhimurium uses the PbgA PD and electrostatic interactions mediated by R215 R216 to negatively regulate LpxC and control LPS abundance during stress. We provide data to support this prediction and demonstrate that the mechanism enhances the ability of *S.* Typhimurium to survive intracellularly in phagocytes, as well as to colonize and kill mice during systemic disease.

## RESULTS

### *S.* Typhimurium uses the periplasmic domain (PD) of PbgA/YejM to negatively regulate LpxC during stress

To determine the contribution of PbgA to LpxC regulation, we compared our wild-type *S.* Typhimurium 14028s genotype and two site-directed deletion-insertion mutants, *pbgAΔ191-586* and *pbgAΔ328-586,* which are deleted for the entire PD or a portion of the globular region for the PD, respectively (**Table 1**) (5, 35). Trans-complementation was achieved by basally expressing PbgA in the *pbgAΔ191-586* mutants from the multi-copy plasmid, pBAD24 (36).

**Table 1.**
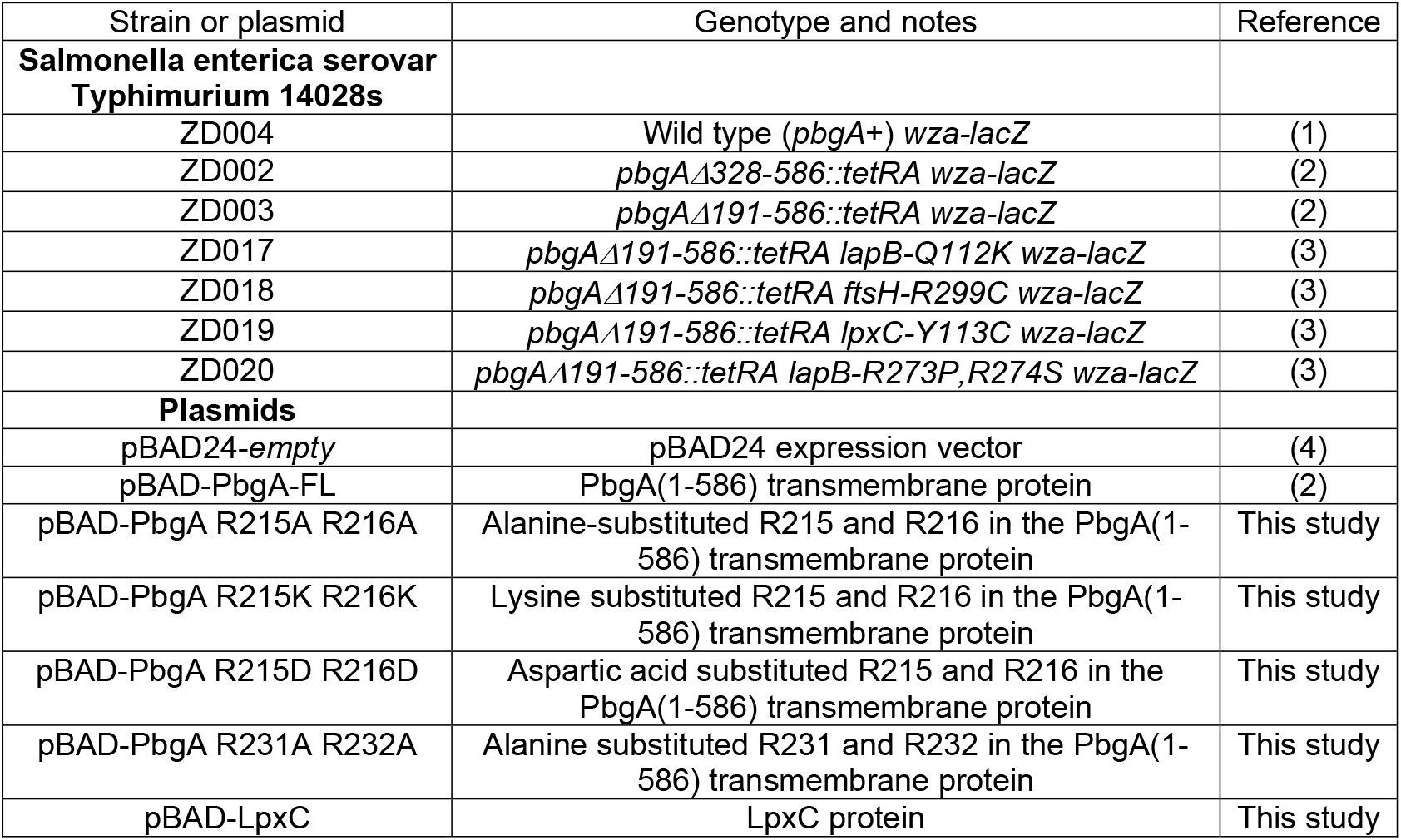
Bacterial strains and plasmids used in this study

*S.* Typhimurium LpxC is a roughly 33-kDa (kD) soluble polypeptide that is localized to the cytosol. During logarithmic growth, LpxC is turned over at a specific rate by proteolysis. During stress, the rate of FtsH activity on LpxC increases causing LpxC and LPS and lipid A-core levels to decrease (37, 38). We focused our attention on the stationary phase of growth (16 hour time point denoted on the curve with a star), **Fig 1A**) for *S.* Typhimurium to test for PbgA-mediated LpxC regulation (**Fig. 1B**). Bacterial lysates were assessed for their LpxC abundance by immunoblotting soluble fractions. Stationary phase wild-type *(pbgA+)* bacteria produced modest but detectable levels of the 33 kD form of LpxC (**Fig. 1B**). Arabinose induction of plasmid-borne LpxC caused a slight over-accumulation of this band relative to the empty vector control genotype, suggesting this is likely LpxC (**Fig. S1**). Consistent with the PD of PbgA promoting LpxC down-regulation, the *pbgAΔ191-586* and *pbgAΔ328-586* mutants accumulated the 33 kD LpxC band in the stationary phase of growth (**Fig 1B**). The LpxC levels for the *pbgAΔ191-586* mutants were consistently greater than for the *pbgAΔ328-586* mutants, suggesting some differential involvement of the PD sub-regions (basic and globular) exists in regards to regulating LpxC concentrations (**Fig. 1B**). The growth rate of *pbgAΔ191-586* mutant *S*. Typhimurium severely contracts near the log-to-stationary phase transition in nutrient-rich broth media (5) (**Fig. 1A**). The differential effects on LpxC may influence the bacterial growth rate since, unlike the *pbgAΔ191-586* mutants the *pbgAΔ328-586* mutants were not similarly attenuated for growth. Transcomplementation of the *pbgAΔ191-586* mutants with PbgA fully decreased the LpxC levels and restored the growth pattern to wild type (**Fig. 1A-B**). In the log phase of growth (OD_600_ = 0.6-0.8), wild type and *pbgA* mutants produced equivalent levels of LpxC under these conditions (**Fig. S2**). Therefore, *S.* Typhimurium requires the PbgA PD to negatively regulate LpxC in response to stationary-phase stress.

**Figure 1.**
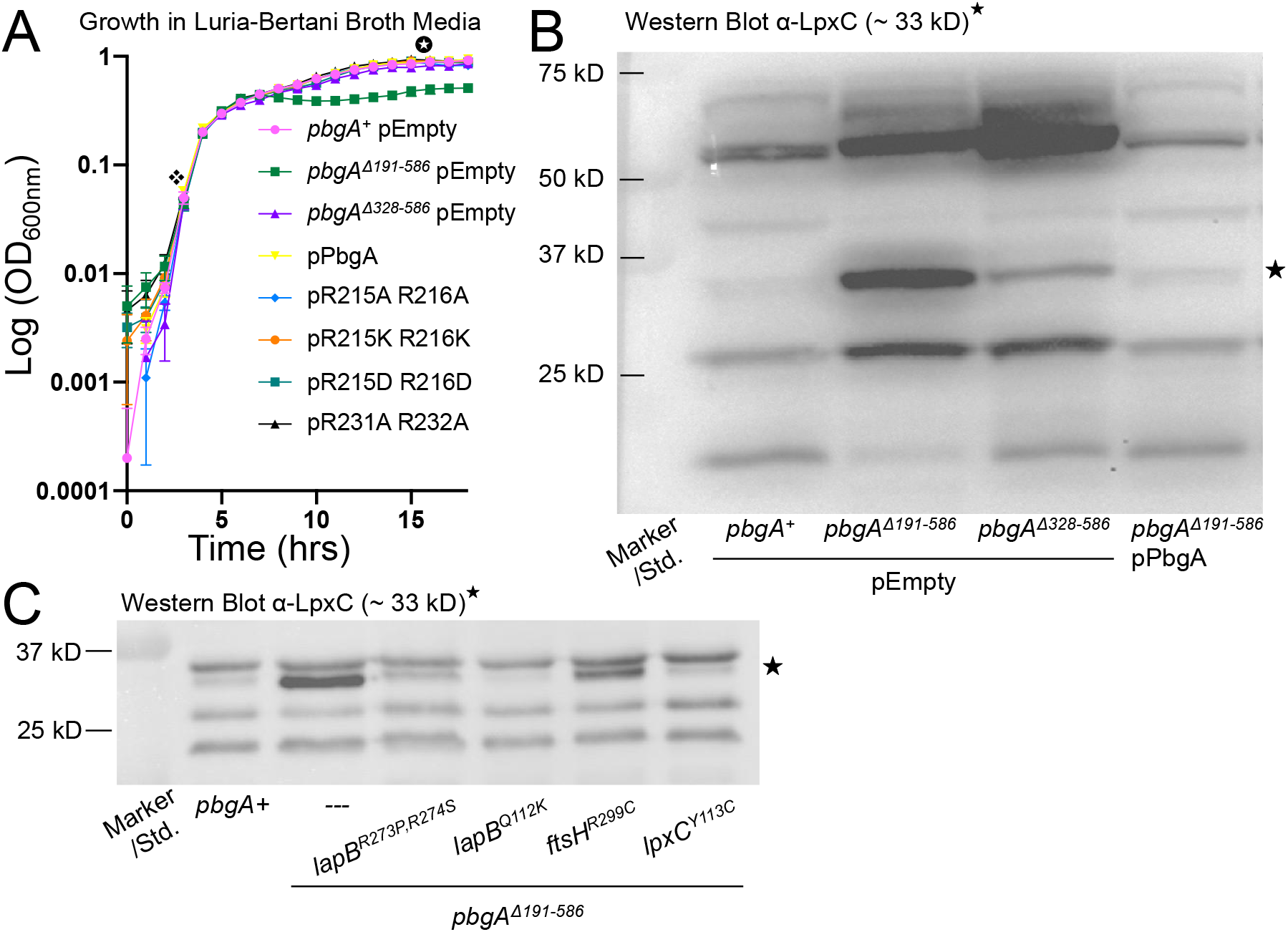
*Salmonella enterica* serovar Typhimurium uses the periplasmic domain (PD) of PbgA (residues 191-586) to negatively regulate LpxC expression in response to stress. (A) Bacterial growth in LB broth. Growth was measured from five biological replicates, each initiated from a single colony re-suspension and conducted as outlined in *materials and methods* section with OD_600_ readings every 15 minutes for 24 hours. Points on the graph represent mean OD_600_ readings at 1-hour intervals and error bars represent the standard error. Asterisks represent the points at which samples were taken for assays. Log phase was defined as OD_600_=0.6-0.8 and stationary phase as 16 hours) (B) Plasmid-bearing *S*. Typhimurium genotypes were grown in 1L of Luria-Bertani (LB) broth supplemented with 100μg/mL ampicillin to the stationary phase of growth. Soluble fractions were concentrated and probed for LpxC abundance. (C) A polyclonal antibody to LpxC (MyBioSource) was used to probe LpxC levels in concentrated soluble fractions isolated from 1L of LB broth. Asterisk denotes the 33kD band that is enriched in *pbgAΔ191-586* mutants.

### Non-synonymous substitutions in LapB, FtsH, and LpxC variably reduce the LpxC levels for the *pbgAΔ191-586* mutants

Our previous suppressor screen yielded *pbgAΔ191-586* isolates with non-synonymous substitutions in LapB/YciM, FtsH, and LpxC (5). The mutations partially restore the growth, lipid A-core accumulation, and virulence defects of the *pbgAΔ191-586* mutants (5). However, we had not determined whether these extragenic mutations influenced the level of LpxC.

After culturing to the stationary phase, wild-type bacteria produced modest levels of the 33 kD polypeptide, while *pbgAΔ191-586* mutants routinely yielded a marked increase (**Fig. 1C**). Relative to the parental mutant genotype, each of the suppressor mutants measured a discernable decrease in the levels of LpxC (5) (**Fig 1C**). Therefore, the suppressive mutations in LapB, FtsH, and LpxC commonly decrease the LpxC levels for the *pbgAΔ191-586* mutants to restore lipid homeostasis and virulence.

***S.* Typhimurium uses the PbgA PD to negatively regulate lipid A-core levels during stress.** To determine whether PbgA-mediated effects on LpxC impacted lipid-A core and LPS biosynthesis, we extracted LPS molecules from identical numbers of viable stationary-phase bacteria and analyzed the relative glycolipid composition by using denaturing gel electrophoresis and staining with ProQ Emerald 300 (5). Lipid A-core (LAC) and short LPS molecules with 1-4 O-antigen repeating units (S1-S4) migrate between 10 and 30 kD and are the best resolved glycolipids in the gels. Our analysis focused on these bands to quantify differences and determine significance (5).

*S.* Typhimurium produces fewer LPS molecules per cell in stationary phase than in log phase growth (**Fig. 2A**) (7, 39, 40). The decrease is partly dependent on the PbgA PD, since *pbgAΔ191-586-mutants* produced qualitatively greater levels of LPS relative to the wild type in stationary phase (**Fig. 2A**). Specifically, the lipid A-core levels for the mutants were significantly four-times greater than the wild type and the transcomplemented genotype (**Fig. 2B)**. The *pbgAΔ328-586* mutants did not accumulate lipid

**Figure 2.**
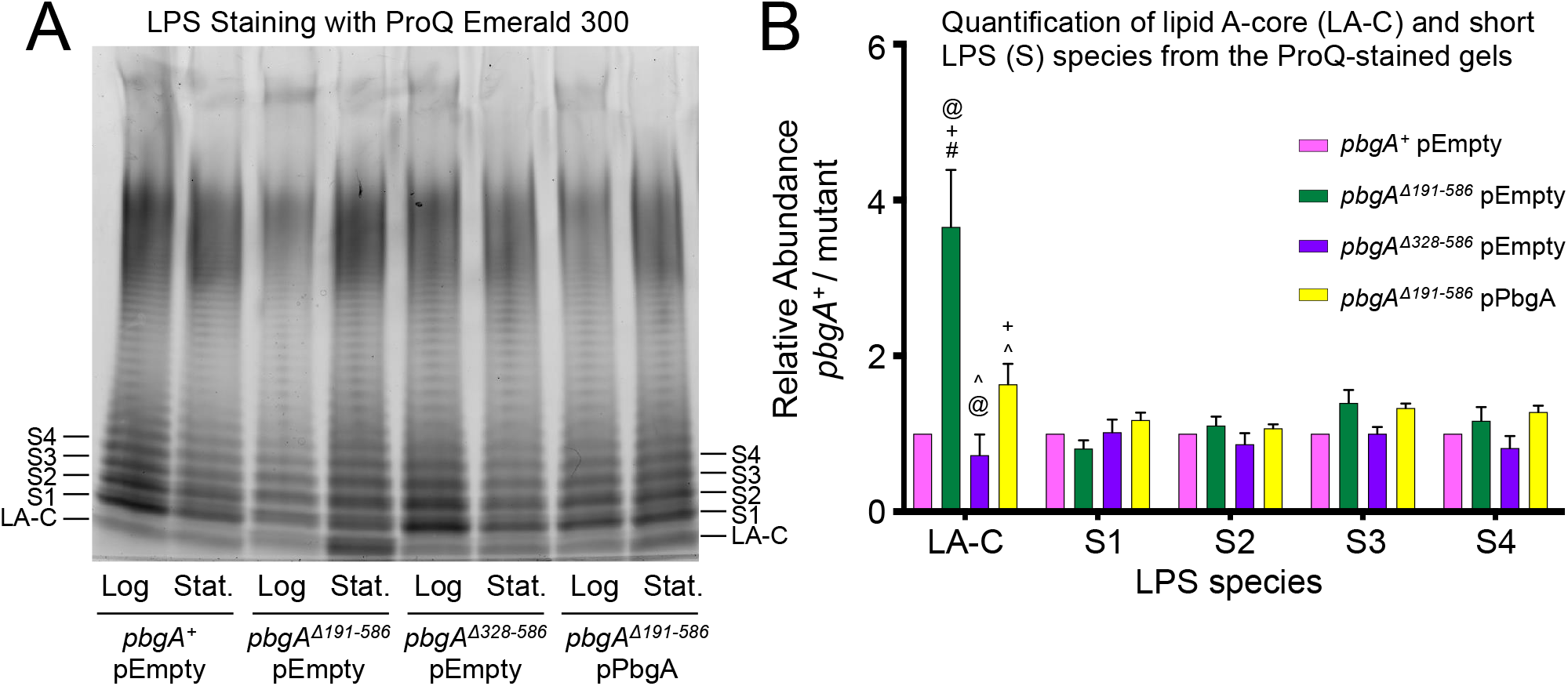
*S*. Typhimurium requires the PbgA PD to decrease lipid A-core levels during stationary phase stress. (A) Each genotype was grown with aeration at 37°C in 5mL of LB broth supplemented with ampicillin (100μg/mL) to the log (OD_600_=0.6-0.8) or the stationary phase. Basal expression from the pBAD promoter without adding inducer or repressor was optimal for genetic transcomplementation and phenotype rescue of the *pbgAΔ191-586* deletion-insertion genotype. LPS was extracted using a hot-phenol method and visualized using ProQ Emerald 300 (Thermo). (B) The levels of the lipid A-core LPS precursor (LA-C) and short LPS species (S1-S4) were quantified for stationary phase LPS using the images and densitometry. Images were obtained using the ChemiDoc™ MP Imaging System (BioRad) and densitometric analysis was performed using BioRad Image Lab^®^ 6.0 Software. All densitometric measurements were normalized to the corresponding band for the wild type strain in the same gel. Three biological replicates were quantified. Values are represented as mean ± standard error. Significance was determined using Tukey’s multiple comparisons (p<0.0001) and is denoted by number symbol (#), up arrow (^), plus sign (+), or the “at” sign (@) to show significance from wild type, *pbgAΔ191-586, pbgAΔ328-586,* or the complemented mutant, respectively. All reached a p-value of <0.0001, with the exception of the comparison between *pbgAΔ328-586* and the complemented mutant, which reached a p-value <0.01.

A-core compared to the wild type despite the routine accumulation of LpxC in these bacteria, albeit to levels that were generally less than the *pbgAΔ191-586* mutants (**Fig. 1B, Fig. 2A-B**) (5). Neither mutant exhibited a significant difference in the amount of short LPS molecules produced (**Fig. 2B**). Therefore, *S*. Typhimurium uses the PbgA PD to negatively regulate LpxC and decrease lipid A-core biogenesis during stress.

***S.* Typhimurium uses the cationic charge of the PbgA R215 R216 side chains, but not the R231 R232 side chains, to negatively regulate LpxC and lipid A-core levels during stress.** The basic region of PbgA is a series of helices and loops that connect the fifth TM segment and the globular region of the PD (**Fig. 3A**) (34, 41). The basic region contains two consecutive helices, each with a pair of dual arginines, R215 R216 and R231 R232. The arginine pairs are highly conserved among enterobacteriaceae (**Fig. S3)** (34, 35, 41). PbgA interacts with the 1-phospho-GlcNAc atoms of lipid A-core via the R216 side-chain atoms, and the R215 and R216 backbone atoms (**Fig. 3B**) (34). The side-chain atoms of R215 form a stabilizing intramolecular contact with the sidechain atoms of residue D192 (34). To test whether the arginine pairs contribute to LpxC regulation, we generated neutral, conservative, and non-conservative charge substitutions at these positions and used a trans-complementation approach.

**Figure 3.**
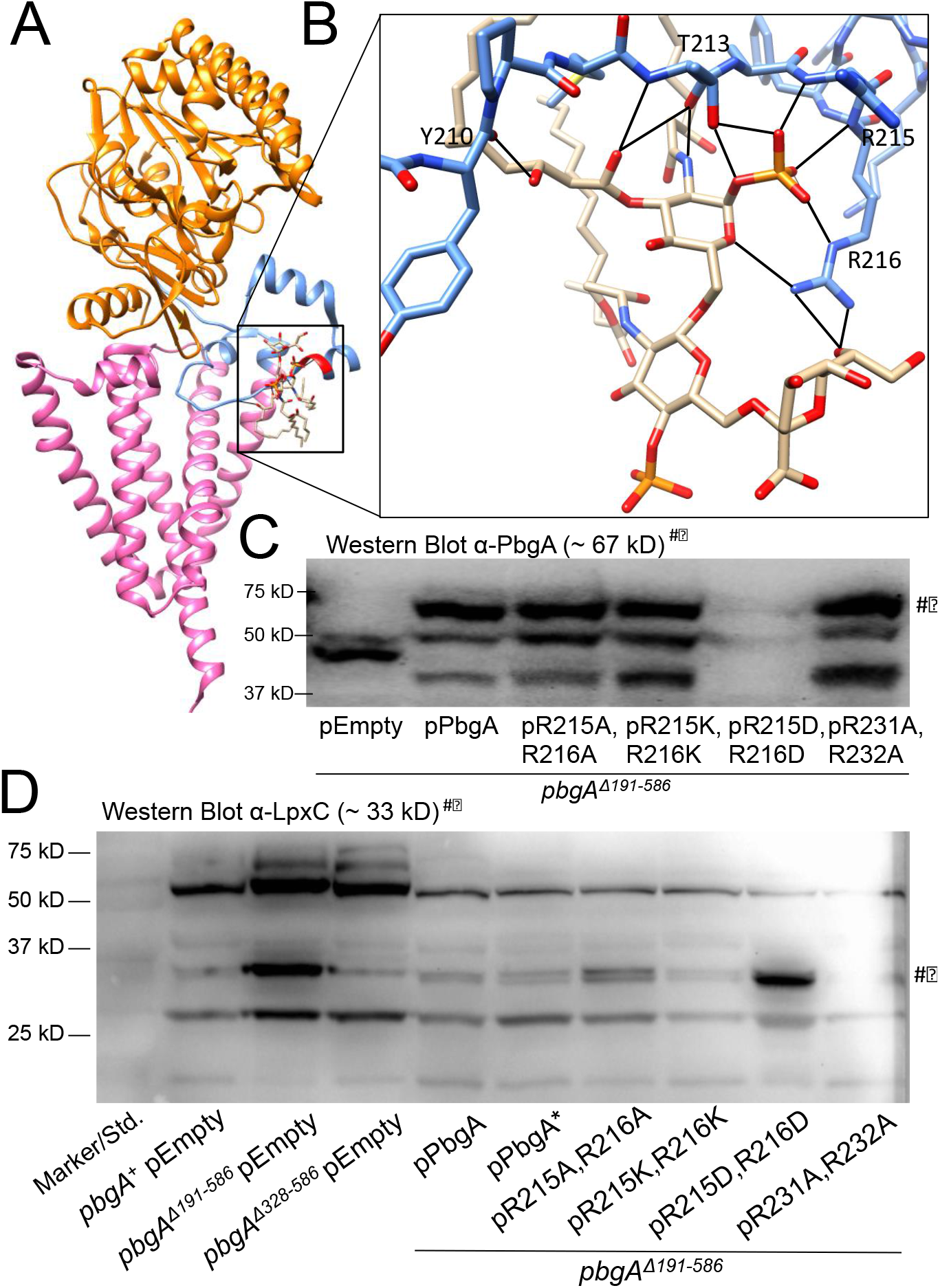
The cationic charge of PbgA R215 R216, but not PbgA R231 R232, is necessary for the negative regulation of LpxC. (A) The three-dimensional crystal structure of PbgA/YejM from S. Typhimurium is depicted (PDB number: 6XLP) and reveals the transmembrane domain (pink), periplasmic basic region (blue), and periplasmic globular region (orange) (34). PbgA R215 R216 is highlighted in red. (B) The inset depicts a close-up of the interactions between the PbgA basic region and 1-phospho-GlcNAc of lipid A-core (tan) at residues Y210 (backbone), T213 (backbone and side chain), R215 (backbone), and R216 (backbone and side chain). Atoms are colored according to charge with the cationic charges in blue and anionic charges in red. (C) PbgA R215A R216A, PbgA R215K R216K, and PbgA R231A R232A are stably expressed in *S*. Typhimurium, but PbgA R215D R216D mutants are possibly unstable. A polyclonal antibody cleared from rabbit antisera raised to purified PbgA^191-586^ polypeptide was used to probe PbgA expression in total membrane fractions isolated from 0.5L of stationary phase broth culture. (D) The cationic charge of R215 R216 is necessary to negatively regulate LpxC. Site-directed amino acid substitution mutagenesis was used to test the role of R215 R216 and R231 R232 in LpxC expression. Both arginine pairs were substituted with neutral alanine residues and the R215 R216 pair were further substituted with conservative and non-conservative charge substitutions. The plasmid-borne alleles were introduced into the *pbgAΔ191-586* deletion-insertion genotype. LpxC levels were probed by immunoblotting soluble fractions of bacteria electrophoresed on an SDS-PAGE gel after growth in 1L broth culture to stationary phase. This blot represents one of six independent experiments. The asterisk represents the addition of 0.02% arabinose, which was used for overexpression (36).

Relative to the *pbgAΔ191-586* mutants expressing the vector, the mutants expressing the PbgA R215A R216A and R231A R232A proteins did not elicit measurable growth defects in broth media and the proteins were stably expressed (**Fig. 1B, Fig. 3C**). The *pbgAΔ191-586* mutants expressing the empty vector produced robust levels of the 33 kD LpxC polypeptide (**Fig. 1B, Fig. 3D)**. Consistent with the first pair of tandem arginines playing a role, the level of LpxC was consistently elevated in the PbgA R215A R216A mutants compared to the wild type and the trans-complemented mutant; however, not to the level of the *pbgAΔ191-586* mutants (**Fig. 3D**). *S*. Typhimurium PbgA R231A R232A mutants produced near wild type LpxC levels (**Fig. 3D**). Therefore, *S*. Typhimurium relies on PbgA R215 R216, but not R231 R232 for stress-induced downregulation of LpxC.

We tested whether the cationic charge of the side chains of PbgA R215 R216 were important for regulating LpxC by substituting for dual lysines or aspartic acids. Like for the wild type and alanine-substituted proteins, the lysine and aspartic acid-substituted variants rescued the stationary-phase growth defect of the *pbgAΔ191-586* mutant strain; however, the aspartic acid protein variants were lowly expressed and possibly unstable in comparison to the wild type and other mutant proteins (**Fig. 1A, Fig. 3C**). The cationic charge of R215 R216 is likely necessary for regulating LpxC, since the lysine-substituted PbgA variants expressed wild-type levels of LpxC in stationary phase (**Fig. 3D**). The *pbgAΔ191-586*-mutant *S*. Typhimurium that expressed the non-conservative charge substitutions, R215D R216D, accumulated LpxC to a greater level than the alanine mutants, but the level was still less than the parental mutant control bacteria, suggesting the attenuated proteins still partly restore the LpxC phenotype of the deletion mutant (**Fig. 3C-D**). These findings support that side-chain interactions mediated by PbgA residues R215 R216 allow *S.* Typhimurium to negatively regulate LpxC in response to stress and that additional residues in the PD are likely involved.

Consistent with the increased abundance of LpxC, the PbgA R215A R216A and R215D R216D mutants produced significantly greater levels of lipid A-core and short-LPS molecules compared to the wild type and the complemented mutant (**Fig. 4A-B**). Consistent with the LpxC levels, the PbgA R215D R216D mutants accumulated greater levels of lipid A-core than the R215A R216A mutants; however, the levels of short-LPS were invariant between the two genotypes (**Fig. 3D, Fig. 4A-B**). Like for LpxC expression, the levels of lipid A-core and short LPS molecules in the PbgA R215K R216K and R231A R232A mutants were statistically the same as the wild type and complemented mutant (**Fig. 3D, Fig. 4A-B**). Therefore, *S*. Typhimurium relies on the cationic charge of the PbgA R215 R216 to negatively regulate LpxC and this allows the bacteria to influence the level of lipid A-core during stress.

**Figure 4.**
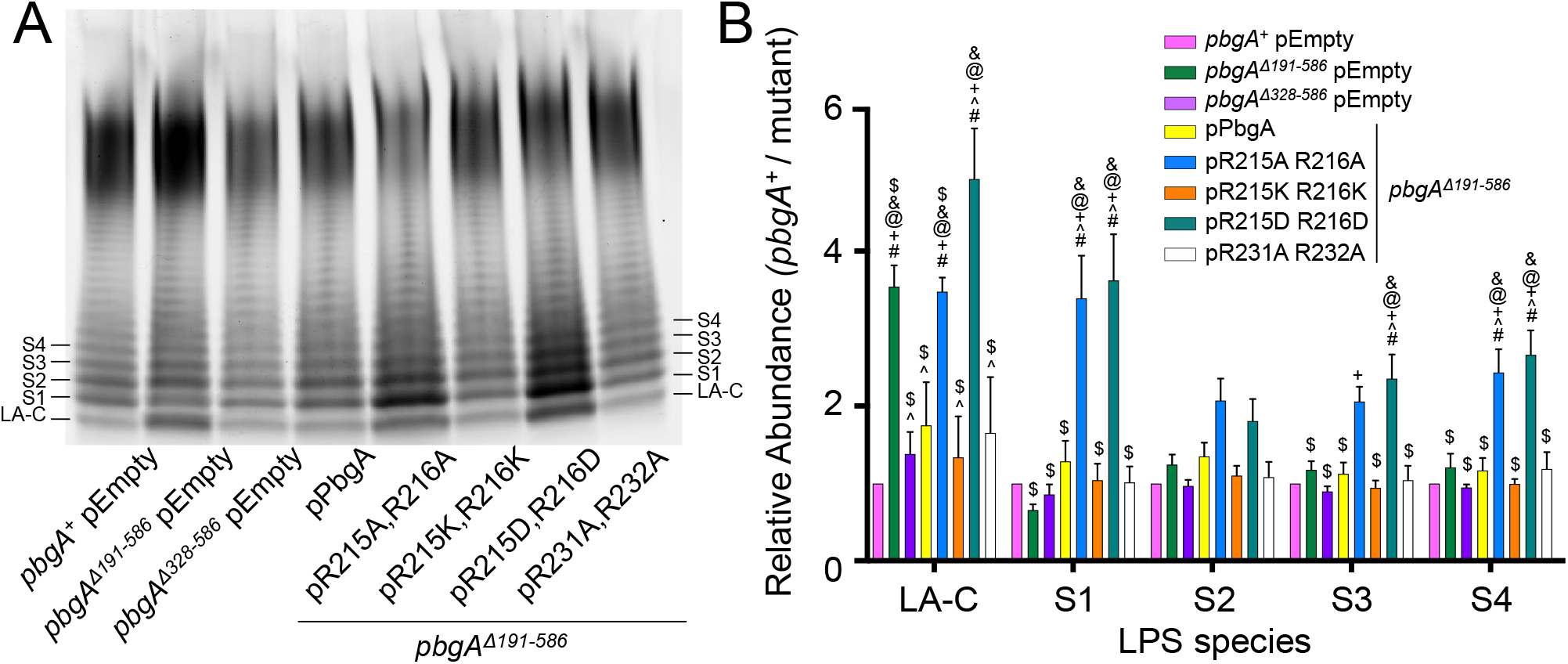
The cationic charge of PbgA R215 R216, but not PbgA R231 R232, is necessary for *S.* Typhimurium to negatively regulate LPS biosynthesis. (A) Strains carrying conservative, non-conservative, and neutral charge substitutions in the arginine pairs described in were grown in 5mL cultures to stationary phase. The LPS molecules were extracted and visualized. (B) The levels of lipid A-core, an O-antigen deficient LPS precursor, and four short O-antigen containing LPS molecules were quantified using densitometry and BioRad Image Lab^®^. Six biological replicates were cultured, extracted, electrophoresed, stained, and quantified to generate the average values depicted here. Data is represented as mean ± standard error. Statistical significance was calculated using Tukey’s multiple comparisons test. Symbols denote significance as follows: #, *pbgA^+^* pEmpty; ^, *pbgAΔ191-586* pEmpty; +, *pbgAΔ328-586* pEmpty; @, *pbgAΔ191-586* pPbgA; &, PbgA R215K R216K; $, PbgA R215D R216D (p<0.0332).

### The cationic charges of PbgA R215 and R216 are necessary for *S*. Typhimurium to survive intracellularly in macrophages

*S.* Typhimurium survives in the endocytic vacuoles of macrophages to cause systemic disease in mice and this requires the PbgA PD (5, 35). To assess the contribution of the dual arginines to intracellular survival, primary bone marrow derived macrophages (BMDMs) from C57Bl/6J mice were infected with the aforementioned genotypes (5). At both time points post infection, the cfu levels of the wild type, the complemented mutant, and the mutants expressing PbgA R215K R216K and R231A R232A reached approximately 10^5^ cfu/mL and did not statistically vary from one another (**Fig. 5**). By contrast, the *pbgAΔ191-586* mutants expressing empty vectors were recovered at 10^4^ cfu/mL less than for the wild type and the complemented mutant at 2 and 6 hours post infection (**Fig. 5**). The *pbgAΔ328-586* mutants expressing empty vectors were recovered at 10^2^ cfu/mL less than wild type **(Fig. 5)**. Salmonellae expressing PbgA R215A R216A and PbgA R215D R216D were partly attenuated and were recovered at 10^1-2^ cfu/mL less than wild type, at both time points (**Fig. 5**). The findings indicate that the cationic charges of the side chains for PbgA R215 R216 are partly necessary for PbgA’s role in enhancing *S.* Typhimurium intracellular survival in macrophages.

**Figure 5.**
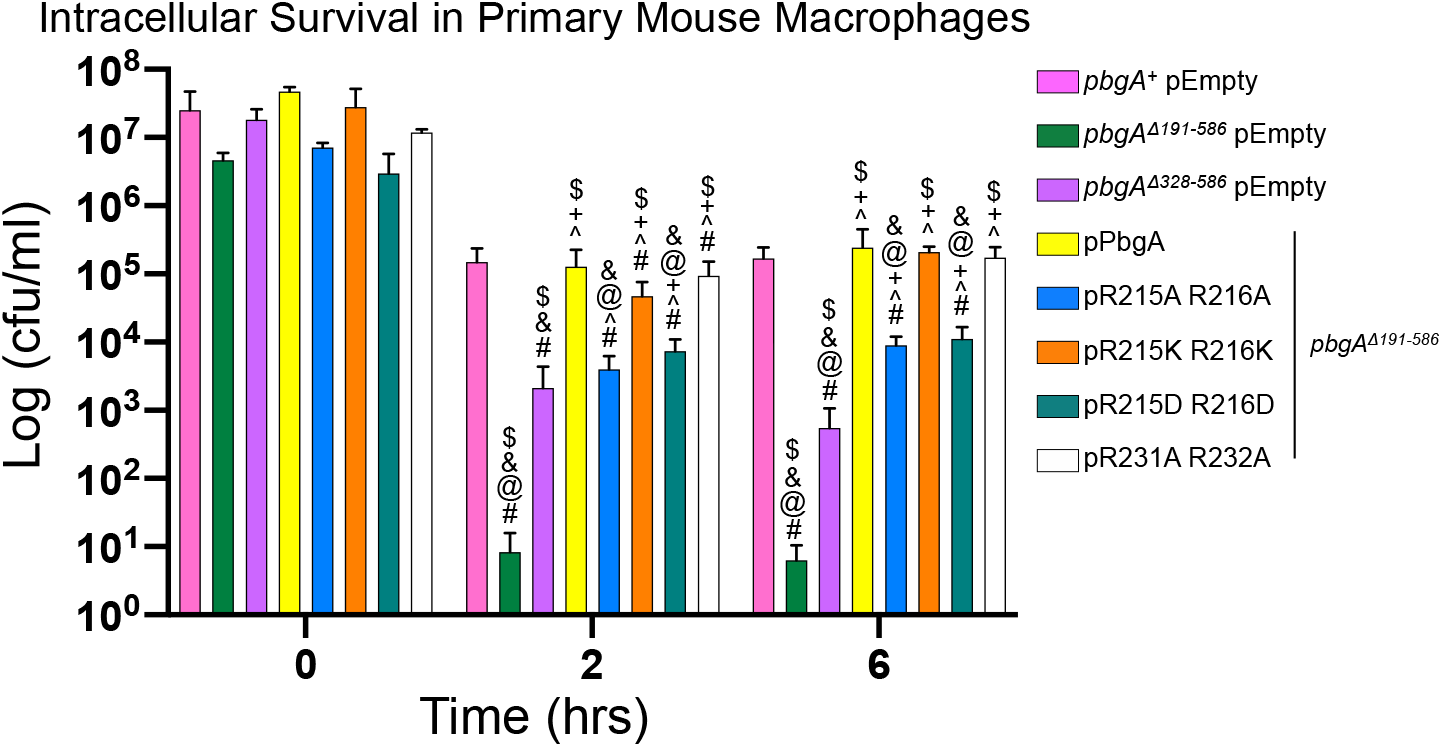
*S*. Typhimurium use the cationic charge of PbgA R215 R216 to survive in primary murine macrophages. Primary bone marrow derived macrophages (BMDMs) from C57Bl/6J mice were infected at a multiplicity of infection of 10:1 with the indicated bacterial strains. Each strain was assessed for intracellular survival in triplicate across three experiments for a total of nine wells at two- and six-hours postinfection. Data is represented as mean ± standard deviation. Significance was determined using Tukey’s multiple comparisons test. Symbols denote significance as follows: #, *pbgA^+^* pEmpty; ^, *pbgAΔ191-586* pEmpty; +, *pbgAΔ328-586* pEmpty; @, *pbgAΔ191-586* pPbgA; &, PbgA R215K R216K; $, PbgA R215D R216D (p<0.0332).

### The side chain charges of PbgA R215 and R216 are necessary for *S.* Typhimurium to colonize the spleens and livers of C57Bl/6J mice

*S.* Typhimurium uses the PD of PbgA to cause systemic disease and lethal bacteremia in mice, so we next tested whether the side-chain charges of the dual arginines were necessary (5). Male and female wild-type C57Bl/6J animals were intraperitoneally inoculated with approximately 10^5^ cfu. At 48 hours, the animals were euthanized and the livers and spleens of the mice were homogenized. Mice infected with the wild type, the complemented mutant, and the mutant expressing PbgA R215K R216K or R231A R232A contained statistically identical numbers, approximately 10^7-9^ cfu/gram of organ tissue (**Fig. 6A-B**). In contrast, the *pbgAΔ191-586* and *pbgAΔ328-586* mutant controls were severely attenuated and only 10^1-2^ cfu were recovered (**Fig. 6A-B**). Mice infected with the PbgA R215A R216A and R215D R216D mutants were colonized with up to 10^2-4^ cfu/gram of organ, which was statistically identical to the *pbgAΔ191-586* mutant controls (**Fig. 6A-B**). Therefore, *S*. Typhimurium uses the cationic charges of PbgA R215 R216 to enhance colonization in mice following intraperitoneal injection.

**Figure 6.**
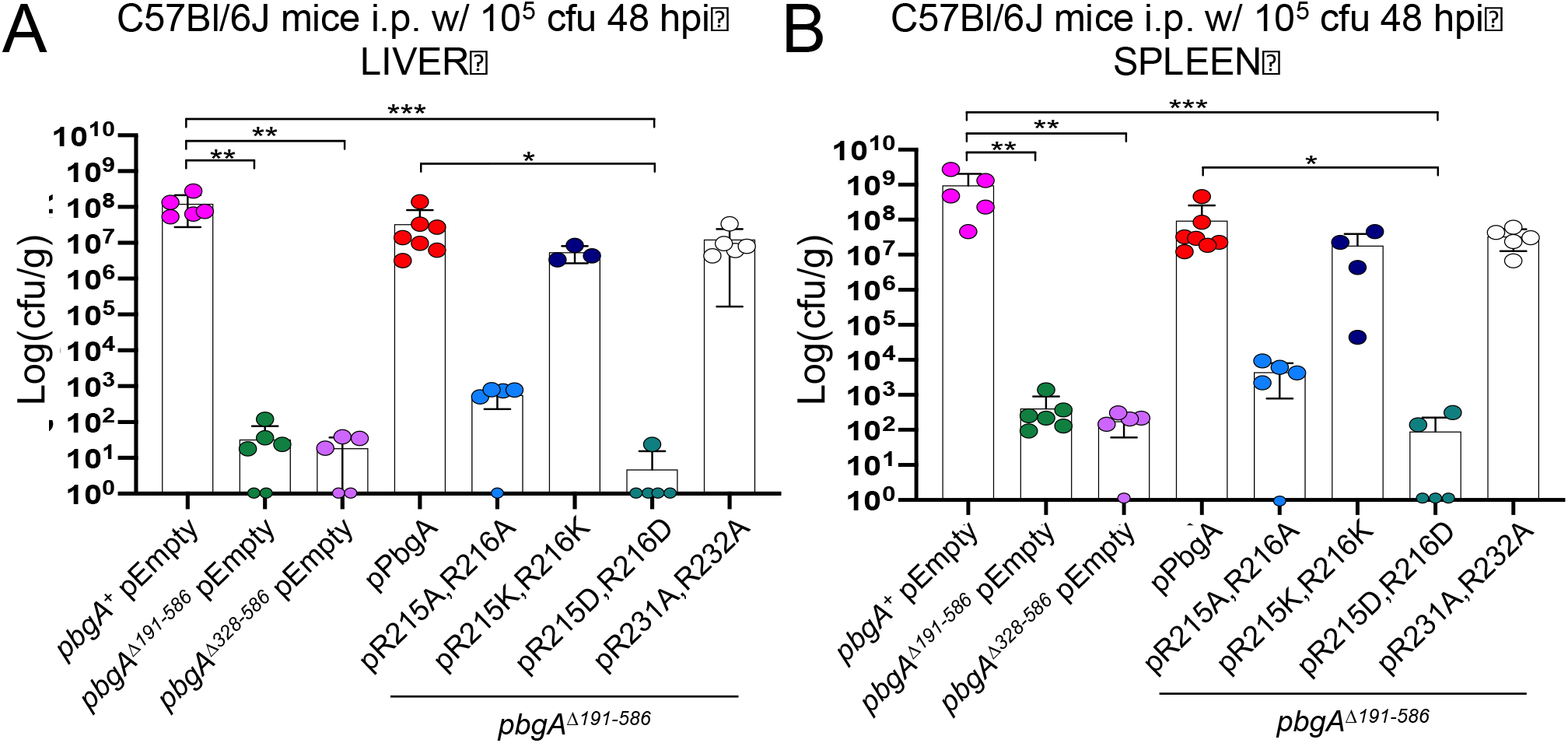
*S.* Typhimurium require electrostatic interactions mediated by the PbgA basic region for colonization of C57Bl/6J mice. Mice were intraperitoneally injected with roughly 10^5^ cfu of one of the indicated bacterial strains suspended in 200μL PBS. Two days later, they were euthanized and their livers (A) and spleens (B) were harvested, homogenized, and plated for CFU. Data is normalized to organ weight and represented as mean ± standard deviation. Statistical significance was calculated using the Kruskal-Wallis test for multiple comparisons (*, p<0.0332, **p<0.0021, ***, p<0.0002).

### *S.* Typhimurium relies on PbgA R215 R216 and R231 R232 to cause lethal bacteremia in C57Bl/6J mice

*S.* Typhimurium *pbgAΔ191-586* suppressor mutants that encode the LpxC^Y113C^ substitution persist in mice without causing lethality (5). The mechanism involves Toll-like receptor 4 (Tlr4), which is the innate immune receptor that binds to the lipid A-core moiety of LPS and controls immune activation and antimicrobial defenses in response to gram-negative bacteria (5, 6). Therefore, we sought to test whether the two pairs of dual arginines were necessary for the ability of *S*. Typhimurium to kill wild type and Tlr4-deficient mice.

Wild-type male or female animals were intraperitoneally infected with 10^3^ cfu and monitored for signs of morbidity and mortality for twenty-one days. Infections with wildtype *S.* Typhimurium were lethal to the wild-type C57Bl/6J mice by 5 days, while the infections with the *pbgAΔ191-586* mutants *pbgAΔ328-586* were not lethal throughout the 21-day duration (**Fig. 7A**). *pbgAΔ191-586* mutants expressing wild-type PbgA and the conservative charge substitutions, R215K R216K, killed all of the wild-type animals by day 8 and 9 respectively, with a mean survival of 7 days post infection (dpi) for both groups (**Fig. 7A).** The mean days of survival did not vary between the groups infected with the complemented mutant and the mutant expressing PbgA R215K R216K, but the mean days of survival for mice infected with these strains was significantly longer than for the group infected with the wild type *S.* Typhimurium (**Fig. 7A**). Therefore, the transcomplementation approach did not fully restore the PbgA-dependent toxicity phenotype in our mouse model, under these conditions. This was consistent in mice lacking Tlr4, which survived an average of 3 days when infected with the wild-type *S*. Typhimurium, but an average of five days for mice infected with the complemented mutants or mutants expressing PbgA R215K R216K (**Fig. 7B**). The *pbgAΔ191-586 and pbgAΔ328-586* mutant infections were not lethal to the Tlr4-deficient animals. Similarly, the infections with the *pbgAΔ191-586* mutants expressing PbgA R215A R216A and R215D R216D were not toxic to the animals under these conditions (**Fig. 7B**). These data are consistent with *S*. Typhimurium relying on the cationic charge of PbgA R215 R216 to colonize and kill mice.

**Figure 7.**
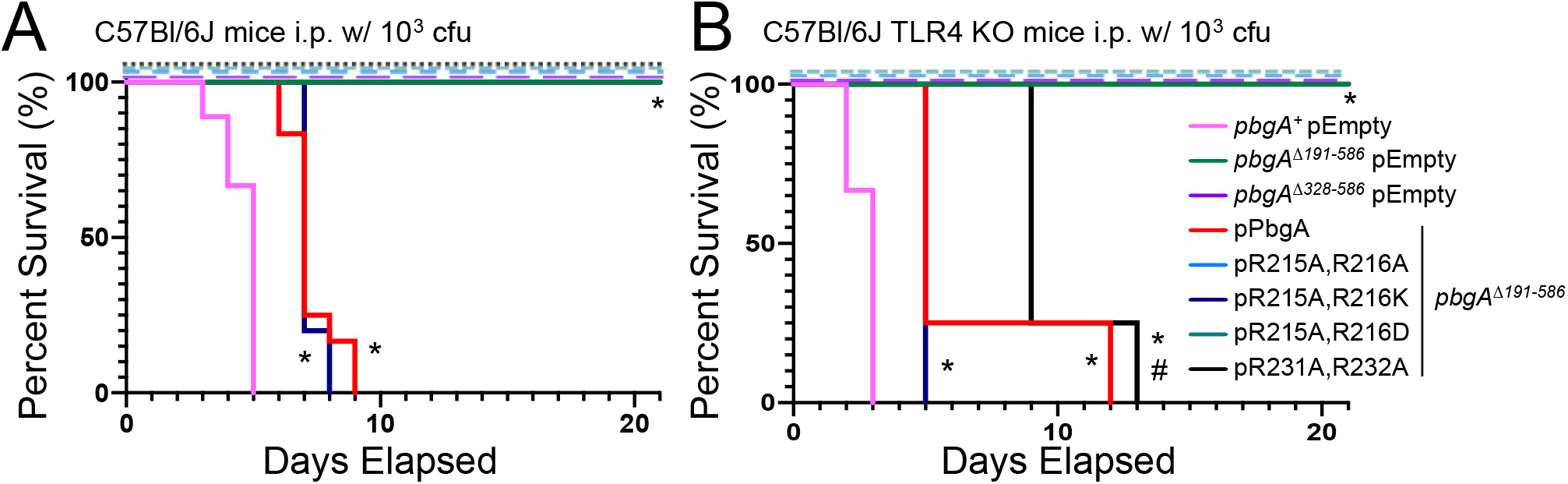
*S*. Typhimurium uses conserved arginine pairs, R215 R216 and R231 R232, to cause lethality in mice. C57Bl/6J (A) and Tlr4 knock out (B) mice were intraperitoneally injected with roughly 10^3^ cfu of one of the indicated bacterial strains suspended in PBS. The wild-type strain, complemented mutant, and PbgA R215K R216K exhibited toxicity towards both strains of mice, which survived to an average of 5, 7, and 7 days, respectively, in the wild-type animals (A) and an average of 3, 7, and 7 days, respectively, for the Tlr4-deficient animals (B). All mice infected with *pbgA^Δ191-586^* pEmpty, *pbgA^Δ328-586^* pEmpty, PbgA R215A R216A, and PbgA R215D R216D survived to the 21-day end point. Tlr4 deficient mice infected with PbgA R231A R232A succumbed to disease by day 13 post-infection, while wild-type animals did not succumb to disease in the 21-day period (B). Significance was calculated using the Log rank (Mantel-Cox) test and symbols indicate statistical significance compared to wildtype, (*, p<0.01; #, p<0.0001).

In the two-day colonization experiments, *S.* Typhimurium *pbgAΔ191-586* mutants expressing PbgA R231A R232A colonized mice to levels that were invariant from the wild type (**Fig. 6A-B**). However, in the lethality assay, the wild-type C57Bl6/J mice infected with PbgA R231A R232A mutants did not perish (**Fig. 7A**). In contrast to the *pbgAΔ191-586* mutants, which were cleared from all infected animals, the mice infected with PbgA R231A R232A still harbored roughly 10^4-5^ salmonellae, indicating that these dual arginines also contribute to PbgA-mediated virulence mechanisms (**Fig. S4**). Tlr4-deficient C57Bl6/J mice that were infected with the PbgA R231A R232A mutants were killed by 13 dpi with a mean survival of 9 dpi, yielding 10^8-9^ salmonellae at the time of death (**Fig. 6D, Fig. 6F**). Therefore, the low dose time-to-death studies suggest that *S.* Typhimurium requires PbgA R231 R232 to cause lethality in mice in a manner that involves Tlr4.

## DISCUSSION

This work supports that *S.* Typhimurium uses the cationic charges of two consecutive arginines for PbgA/YejM to negatively regulate lipid A-core biosynthesis in response to stress. The sensory and signal transduction mechanism likely involves contributions from the IM-anchored cytosolic protein, LapB/YciM, and the integral IM protease FtsH, and the ability of LapB-FtsH to negatively regulate the level of LpxC in the cytosol (**Fig. S5**). PbgA’s contribution to regulating LpxC is critical for *S.* Typhimurium to survive in macrophages and to cause systemic disease mice (**Fig. 5; Fig. 6**). The need for *S*. Typhimurium to down-regulate lipid A-core biogenesis during intracellular survival and systemic pathogenesis could have multiple biological purposes.

In the resource allocation model, the ability to negatively regulate LpxC and lipid A-core biogenesis serves as a mechanism by which enterobacteriaceae free up fatty acid and N-acetyl glucosamine substrates for use in other metabolic pathways (30, 39, 40, 42, 43). Instead of a strategy to preserve key metabolites, enterobacterial downregulation of lipid A-core biosynthesis might alternatively serve an offensive purpose (6). The bilayer couple model of outer membrane vesicle (OMV) formation predicts that increasing the level of an amphipathic molecule in the OM outer leaflet causes the outer leaflet to expand relative to the inner leaflet. Insertion and expansion of the outer leaflet induces curvature and vesicle formation (44). Perhaps conditions in host environments elicit *S.* Typhimurium to produce more GPLs on the outer leaflet and fewer lipid A-core and LPS molecules. We have established that *S*. Typhimurium constitutively invert GPLs into the outer leaflet of the OM where they become substrates for lipid A-core and phosphatidylglycerol acylation by the PagP enzyme (45, 46). Stress-induced decreases in LPS could be coordinated with increased phospholipid inversion to regulate vesicle formation in the acidic environment of phagocyte endosomes. By this rationale, the periplasmic arginines for PbgA/YejM may sense lipid anions, such as lipid A-core, that accumulate in the IM to regulate vesicle formation under stressful conditions that *S*. Typhimurium encounter in the host.

High-resolution crystal structures of PbgA reveal contacts between lipid A-core and the backbone of R215, as well as the side chain of R216 (**Fig. 3B**) (34, 41). The R215 side chain also forms a contact with the D192 side chain, which is positioned near the C-terminal end of fifth TM segment (34). Since salmonellae expressing PbgA with dual lysine substitutions are not defective in regulating LpxC, surviving in macrophages, or causing systemic pathogenesis in mice, but salmonellae with dual alanine substitutions are defective, the data support that electrostatic side-chain interactions between R216 and lipid A-core, and between R215 and D192 contribute to PbgA’s ability to regulate LpxC.

The prevailing data support that PbgA R215 R216 participate in electrostatic interactions that are necessary for *S*. Typhimurium to down-regulate LpxC and decrease lipid A-core biosynthesis in response to stress. Biochemical studies in *E. coli* indicate that PbgA binds LapB through the TM regions of each protein (31, 34). *E. coli* LapB binds FtsH and LpxC, but the residues that are necessary are are not known (27). *E. coli* PbgA R215A R216A mutant proteins maintain interactions with LapB, so we predict that PbgA-LapB binding is not directly impacted by the interactions of R215 and R216 (34).

In our model, we hypothesize in non-hazardous and replete environments, *S.* Typhimurium PbgA binds LapB and holds it in an inactive conformation (**Fig. S5**). During stress, an anionic lipid molecule, perhaps lipid A-core, accumulates at the periplasmic leaflet of the IM and binds PbgA R215 R216. PbgA-lipid A-core binding causes a putative conformational change in the PbgA-LapB complex, which alters the proteolytic activity of FtsH on LpxC (**Fig. S5**). Unlike the wild type, the R215A R216A, and R215K R216K mutant proteins, the R215D R216D mutant proteins are lowly expressed and possibly unstable compared to the wild type and other mutant proteins (**Fig. 3C**). Like the dual alanine mutants, the dual aspartic acid mutants are highly defective at restoring the virulence defects of the parental mutant genotype, but are capable of restoring the growth defect of the parental mutant genotype in broth culture (**Fig. 1A)**. Since the lysine-substituted proteins are not defective for regulating LpxC or lipid A-core, and are as virulent as the wild type control, our results indicate that the cationic charge of the side chains of these residues likely enables PbgA to regulate LpxC (**Fig. 3D, Fig. 4–7**).

During structural analysis, a second pair of consecutive arginines at positions 231 and 232 attracted our attention (**Fig. S3**) (35). We predicted that these residues might similarly contribute to *S*. Typhimurium LpxC regulation. However, the results suggest that these arginines are not required for salmonellae to regulate LpxC and lipid A-core during stress under the *in vitro* conditions we tested here, nor are they necessary for *S*. Typhimurium to survive in mouse macrophages, nor mouse spleens nor livers after intraperitoneal injection (**Figs. 3C; Fig. 4–6**). In contrast, the mouse lethality studies support that PbgA R231A R232A mutants are not toxic to wild type animals at low doses, and that the mutant salmonellae persist at moderate titers in the spleens and livers for at least 21 days (**Fig. 7A)**. This persistence phenotype resembled that of the *pbgAΔ191-586 lpxC^Y113C^* suppressor mutant (5). Like for the suppressor genotype, the PbgA R231A R232A infections were lethal to Tlr4-deficient animals with a mean time-to-death of 14 days for the mutant compared to 3 days for the wild type control (**Fig. 7B**). Collectively, these results suggest that *S.* Typhimurium uses PbgA to influence the host-immune response to lipid A-core, and to enhance virulence by multiple mechanisms. Future biochemical and phenotypic assays will deduce the exact role of PbgA R231 R232 in *S.* Typhimurium pathogenesis.

Our results continue to support a critical role for the PbgA PD in regulating OM lipid homeostasis and disease pathogenesis in *S*. Typhimurium. The recent surge of experimental attention on PbgA/YejM in proteobacteria indicates that knowledge of the biochemical mechanisms for PbgA-LapB and LapB-FtsH mediated LpxC regulation will be critical to understanding antimicrobial resistance and the immune response to enteric pathogens.

## METHODS

### Ethics statement

All animal procedures were carried out with approval from the University of Oklahoma Health Sciences Center Institutional Animal Care and Use Committee under protocol number 19-015-ACI. The procedures used in this study strictly adhered to the guidelines found in the National Research Council’s Guide for the Care and Use of Laboratory Animals (National Research Council. 2011. Guide for the Care and Use of Laboratory Animals, 8^th^ ed. National Academies Press, Washington DC.)

### Bacterial Strains and Culturing Conditions

The bacterial strains used in this study were all derivatives of the *Salmonella enterica* serovar Typhimurium genotype 14028s, which contains a chromosomally-integrated *wza-lacZ* gene promoter fusion (Table 1) (47). Suppressors of *pbgAΔ191-586::tetRA* were isolated in a previously published suppressor screen and streaked onto Luria-Bertani agar plates containing the LacZ indicator substrate, 5-Bromo-4-Cloro-3-Indolyl ß-D-Galactopyranoside (X-gal) at a concentration of 20μg/mL (5). All other strains in this study contain the pBAD24 plasmid, which was either left empty or contained the full-length PbgA protein (35, 36). Each plasmid-bearing strain was streaked onto Luria-Bertani agar plates containing X-gal at a concentration of 20μg/mL and 100μg/mL ampicillin to maintain the pBAD24 plasmid. The bacteria were isolated from −80°C glycerol stocks, weekly. Cultures were routinely started with a single colony inoculated into LB-broth medium and shaken, or rotated at 250 revolutions per minute, aerobically at 37°C. Ampicillin was added to the growth medium to maintain the plxasmids. Log phase was defined as an optical density at 600nm (OD_600_) of 0.6 to 0.8, and stationary phase growth was determined to be 16h post single-colony inoculation (**Fig. 1A**) (5). The complementation genotype contains the full-length protein basally expressed from the pBAD24 plasmid in the *pbgAΔ191-586* mutant strain background. Arabinose was used for overexpression (36).

### Genetics

The *pbgAΔ191-586::tetRA* and *pbgAΔ328-586::tetRA* insertion-deletion mutants were generated by methods previously described (35). Point mutants were generated using overlapping PCR primers, both containing the desired mutations (**Table 2**). The template for the PCR reaction was purified pBAD24-PbgA, which was generated by methods previously described (35). AccuPrime™ *Pfx* DNA polymerase (Thermo) and the corresponding buffer were used in 50μL reactions. PCR products were Dpn1 treated for one hour at 37°C and isolated using the GeneJET PCR Purification Kit (Thermo). The DNA was then transformed into DH5α and single colonies of the transformants were grown overnight. Plasmids were purified using the GeneJET Plasmid Miniprep Kit (Thermo) and mutations were confirmed by sequencing. The mutated plasmids were then electroporated into competent *pbgAΔ191-586::tetRA* cells.

**Table 2.**
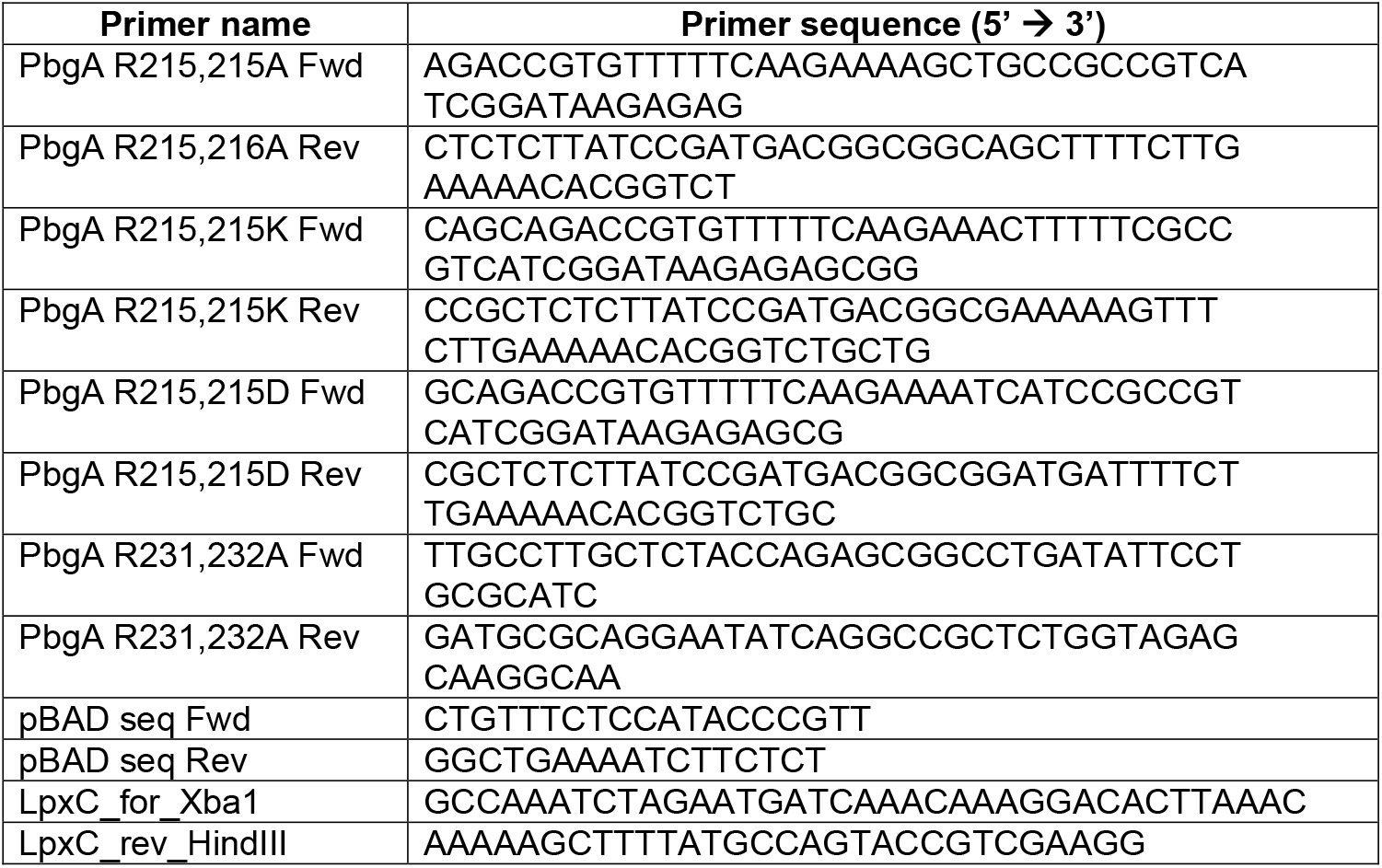
Primers used in this study

### Clearing of PbgA antisera

Anti-PbgA antibodies were cleared from rabbit antisera as previously described (35). 5mL of rabbit antisera was eluted over a protein-A column (Thermo) to isolate Fc/Fab fragments. Non-specific *S*. Typhimurium cross-reacting antibodies were cleared by incubating 50μL of the Fc/Fab elution with 50μL of bacterial cell lysate of *pbgAΔ191-586::tetRA* mutant *S*. Typhimurium in 50mM Tris-HCl pH 8.0 10mM EDTA and 5% non-fat dried milk for 4 hours at room temperature. We routinely observed a contaminating band in the deletion mutant that was roughly 40 kD. The antisera were raised against PbgA191-586 peptides and the remaining TM coding sequence on the genome of *pbgAΔ191-586::tetRA* is predicted to be ~ 22.5 kD.

### Western Blotting

For all western blots, bacteria was grown in 0.5L, for PbgA blots, or 1L, for LpxC blots, of LB broth supplemented with 100μL/mL ampicillin. Bradford assays were used to measure the protein concentration of total membrane fractions (PbgA blots) or soluble fractions (LpxC blots). 20μg of protein was loaded onto a 12% SDS-PAGE gel, electrophoresed, and transferred onto a polyvinylidene fluoride (PVDF) membrane using the Mini Trans-Blot Cell (BioRad) apparatus for wet transfer at 100 volts for 50 minutes. The membrane was washed in Tris-buffer saline with tween 20

(TBST) and blocked overnight at 4°C in 5% non-fat dried milk in TBST. For PbgA, the primary antibody was diluted 1:250 in TBST and applied to the blocked membrane and incubated at room temperature for 2 hours. For LpxC, the primary antibody (MyBioSource) was diluted 1:10,000 in TBST, applied to the blocked membrane, and incubated at room temperature for 1 hour. For both, PbgA and LpxC blots, the anti-rabbit-HRP secondary antibody (Cell Signaling) was diluted 1:5000 in TBST and incubated for 1 hour at room temperature. Blots were imaged after detection with Amersham ECL Prime Western Blotting Detection reagent (GE Healthcare) using the BioRad ChemiDoc MP Imager.

### Growth Curve

Growth curves were generated in a Bioscreen C growth curve analyzer. A single colony was resuspended in 180μL LB broth with ampicillin (100μg/mL) and serially diluted to 10^-3^ to assess growth patterns of the deletion mutants and point mutants. Bacteria were incubated with continuous agitation and OD_600_ was measured and recorded every 15 minutes. Five biological replicates were performed for each bacterial strain and the results reflect the average.

### LPS extraction and visualization

This method was slightly modified from Cian et al, 2019 (5). Bacteria were cultured in 5mL Luria-Bertani Broth (LB broth) supplemented with 100μg/mL ampicillin. Each strain was normalized to an OD_600_ of 2.5, spun down in a 1.5mL microcentrifuge tube, and resuspended in 200μL of sterile water. To confirm that we were extracting from the same number of viable bacteria, 20μL of the 200μL resuspension was plated on LB-agar plates containing ampicillin (100μg/mL) and the cfu/mL values were compared and shown to be statistically identical across experiments. 2μL of 2% SDS were added to the remaining 180μL, which was then incubated in a boiling water bath for ten minutes. After cooling, 5μL of proteinase K (New England BioLabs) was added to each sample and the samples were incubated overnight in a 59°C water bath. 182μL hot phenol was added to each sample and the samples were incubated at 68°C for 10 minutes and immediately transferred to an ice water bath for an additional 10 minutes. The samples were spun down in a temperature-controlled microcentrifuge at 4°C and 4500 rpm for 10 minutes and immediately placed back in the ice water bath. The top LPS-containing aqueous layer was transferred to a new 1.5mL microcentrifuge tube. For visualization, samples were combined with 4x Laemmli Sample Buffer (BioRad) and loaded into a 4-20% SDS-PAGE gradient gel (Mini-PROTEAN^®^ TGX™ Precast Gel; BioRad). The gel was stained according to the ProQ Emerald 300 lipopolysaccharide staining kit (Thermo) protocol.

### Murine macrophage infections

Primary bone marrow-derived murine macrophages (BMDMs) were prepared by harvesting the marrow from the femurs of 6- to 8-week old C57Bl/6J mice bred in-house (5). Macrophages were seeded at 2.5×10^5^ cells per well and infected at a multiplicity of infection of 10. Infected macrophages were incubated at 37°C under 5%CO2 for 1h. The infected cells were washed and aspirated three times with phosphate-buffered saline (PBS) to remove extracellular bacteria. RPMI+FBS with 100μg/ml of gentamycin was added to kill remaining extracellular bacteria. Infected cells were incubated for an additional 1h at 37°C under 5%CO2. At 2h post-infection (hpi), PBS+0.1% Triton was added for lysis and monolayers were gently scraped and collected with a pipette. Three wells per bacterial genotype were assessed per time point. Surviving intracellular colony-forming units (cfu) were enumerated by plating serial dilutions in PBS. After 2 hpi, the wells for the 6hpi time point were aspirated and RPMI+FBS containing 10μg/ml of gentamycin was added to kill bacteria that became extracellular during infection. At 6hpi, macrophages were lysed and surviving intracellular cfu were enumerated.

### Mouse infections

Male and female C57Bl/6J mice were purchased from The Jackson Laboratory and bred in-house under pathogen-free conditions. To measure the ability of *S.* Typhimurium to survive systemically and colonize the spleens and livers of mice, 6-to 8-week old mice were intraperitoneally infected with roughly 5×10^5^ cfu diluted in PBS. At 48 hours, the mice were euthanized and the livers and spleens were dissected, weighed, and homogenized in PBS-0.1% triton X-100. They were then serially diluted and plated on LB-agar plates supplemented with X-gal (20μg/mL) and ampicillin (100μg/mg) to enumerate colony-forming units.

For time-to-death assays, mice were intraperitoneally inoculated with 5×10^3^ CFU diluted in PBS. Mice were sacrificed either at death, defined by physical signs of distress and weight loss, or at the twenty-one-day end point. CFU were enumerated as outlined for the two-day infection.

### Statistical analysis

All statistical analyses were performed and graphs were prepared using GraphPad Prism (version 8; GraphPad Software, La Jolla, CA, USA).

## Acknowledgements

We would like to acknowledge the other members of the Dalebroux Lab including Meli Cian, Keaton Minor, and Aaron Zahn who provided experimental and technical advice throughout the project. This work was funded by grant 5T32AI007633-17, awarded to Nicole P. Giordano, and grants P20GM10344 and R01AI139248, awarded to Z. D. Dalebroux.

